# An Out-of-Patagonia dispersal explains most of the worldwide genetic distribution in *Saccharomyces eubayanus*

**DOI:** 10.1101/709253

**Authors:** Roberto F. Nespolo, Carlos A. Villarroel, Christian I. Oporto, Sebastián M. Tapia, Franco Vega, Kamila Urbina, Matteo De Chiara, Simone Mozzachiodi, Ekaterina Mikhalev, Dawn Thompson, Pablo Saenz-Agudelo, Gianni Liti, Francisco A. Cubillos

## Abstract

*Saccharomyces eubayanus* represents missing cryotolerant ancestor of lager yeast hybrid and can be found in Patagonia in association with *Nothofagus* forests. The limited number of isolates and associated genomes available has prevented to resolve the *S. eubayanus* origin and evolution. Here, we present a sampling effort at an unprecedented scale and report the isolation of 160 strains from ten sampling sites along 2,000 km distance in South America. We sequenced the genome of 82 strains and, together with other 25 available genomes, performed comprehensive phylogenetic analysis. Our results revealed the presence of three main Patagonia-B lineages together with dozens of admixed strains distributed in three mosaic clusters. The PB-1 lineage isolated from Tierra del Fuego exhibited the highest genetic diversity, lowest LD blocks and highest *F*is values compared to the other lineages, suggesting a successful adaptation to cold temperatures in extreme environments and greater inbreeding rates in Tierra del Fuego. Differences between lineages and strains were found in terms of aneuploidy and pangenome content, evidencing a lateral gene transfer event in PB-2 strains from an unknown donor species. Overall, the Patagonian lineages, particularly southern populations, showed a greater global genetic diversity compared to Holarctic and Chinese lineages, supporting the scenario of a *S. eubayanus* colonization from Patagonia and then spread towards northern and western regions, including the Holarctic (North America and China) and New Zealand. Interestingly, fermentative capacity and maltose consumption resulted negatively correlated with latitude, indicating a better fermentative performance in norther populations. Our genome analysis together with previous reports in the sister species *S. uvarum* strongly suggests that the *S. eubayanus* ancestor could have originated in Patagonia or the Southern Hemisphere, rather than China, yet further studies are needed to resolve this conflicting scenario. Understanding *S. eubayanus* evolutionary history is crucial to resolve the unknown origin of the lager yeast and might open new avenues for biotechnological applications.

## INTRODUCTION

There are at least 1,500 species of yeasts, which can be found on a broad range of substrates including fruit skin, cacti exudates and soil, where they can be either inert or pathogenic (Guz 2011). Some species from the Saccharomycotina subphylum have developed the ability to ferment simple sugars from fruits to produce alcohol. In this way, fermentation became a key innovation that led to the diversification of fermentative yeasts about 100 million years ago (MYA), coinciding with the appearance of Angiosperms (Piskur et al. 2006; Dashko et al. 2014). The monophyletic *Saccharomyces* genus is currently composed of eight distinct species (Dujon and Louis 2017), including the partially domesticated *S. cerevisiae* and other non-domesticated species, such as *S. eubayanus* (Borneman and Pretorius 2015). This clade contains some of the most important species involved in alcohol and bread fermentation, likely due to their ability to grow in the absence of oxygen (anaerobic fermentation)(Hagman et al. 2013). Given the economic importance of this clade, as well as the wealth of genomic information that has been produced in the past decade, particularly for the model organism *S. cerevisiae* (Liti et al. 2009; Schacherer et al. 2009; Gallone et al. 2016; Goncalves et al. 2016; Legras et al. 2018; Peter et al. 2018), natural populations of *Saccharomyces* are excellent models for understanding genome evolution and adaptation in the wild.

*S. cerevisiae* was the first sequenced eukaryote, and recently the large amount of isolates in this species and associated genomic data provide exceptional new insights into the genomic processes that drive environmental adaptation and genome evolution between isolates (Yue et al. 2017; Legras et al. 2018; Peter et al. 2018). Given the feasibility to rapidly and cost-effectively sequence full genomes, other *Saccharomyces* genomes have been fully obtained (Dujon and Louis 2017). *Saccharomyces* species harbour different genetic structures, population histories and unique phenotypic properties. Despite these advances, the number of isolates for which both fully annotated genomes and phenotypic data are available is still low. In many cases, only a handful of isolates from a species have been studied and therefore the identification of genomic features responsible for local adaptation and evolutionary changes is less well documented compared to *S. cerevisaie*, limiting the understanding of adaptation processes in the genus.

In nature, several *Saccharomyces* inter-species hybrids have been found. An example of this includes the workhorse of the modern brewing industry, *S. pastorianus*, a hybrid between *S.* c*erevisiae* and the cold-tolerant *S. eubayanus* (Baker et al. 2015; Krogerus et al. 2017). Despite the industrial importance of *S. pastorianus*, much of the natural history of this hybrid remains obscure, largely because the *S. eubayanus* parental strain was only recently isolated (Libkind et al. 2011). The combination of precise alleles gives the hybrid *S. pastorianus* a series of competitive advantages in the fermentative environment. For example, efficient maltotriose utilization was inherited from *S. cerevisiae*, while fermentation at low temperatures and maltose utilization is the legacy of the cryotolerant *S. eubayanus* (Hebly et al. 2015; Krogerus et al. 2017; Brickwedde et al. 2018; Eizaguirre et al. 2018; Baker et al. 2019). Providing a unique fermentation profile for brewing, *S. eubayanus* can efficiently grow at a lower range of temperatures (4°C – 25°C) compared to *S. cerevisiae*, however the genetic basis of this advantage is yet unknown (Baker et al. 2015). *S. eubayanus* was originally isolated from *Nothofagus* trees in the Argentinian Patagonia (Libkind et al. 2011) and since then it has been isolated in New Zealand (Gayevskiy and Goddard 2016), North America (Peris et al. 2014) and East Asia (Bing et al. 2014). However, the evolutionary origin of *S. eubayanus* is still controversial. While this species has been isolated from South American *Nothofagus* trees recurrently (Eizaguirre et al. 2018) and only a handful of isolates have been recovered from trees in China and North America (Bing et al. 2014; Peris et al. 2016), a subset of the strains from China have been reported as the earliest diverging lineage, suggesting an Asian origin of the species (Bing et al. 2014), although these findings have been challenged (Eizaguirre et al. 2018).

Molecular profiling indicates that *S. eubayanus* is composed of three populations, besides the early diverging lineage of West China. These populations include a ‘Holarctic’ cluster (a group of related strains from Tibet and North America) and two Patagonian populations denominated: ‘Patagonia A’ and ‘Patagonia B’ (Peris et al. 2016). The populations, can be further divided into six subpopulations, denominated: PB-1, PB-2, PB-3, Holarctic, PA-1 & PA-2 (Peris et al. 2016; Eizaguirre et al. 2018). Whole genome sequence comparison among wild *S. eubayanus* strains indicate that, thus far, the Holarctic lineage is the closest relative of the lager yeast (Peris et al. 2016; Eizaguirre et al. 2018). Interestingly, multi-locus sequence comparisons have indicated that the nucleotide diversity of *S. eubayanus* Patagonian populations is greater than that of the West China and the Holarctic (North American) lineages, suggesting a greater colonization success in South America (Peris et al. 2016). However, until now only a handful of *S. eubayanus* genomes per population have been fully sequenced preventing a detailed population genomics portrait.

Here, we isolated 160 *S. eubayanus* strains from bark samples obtained from *Nothofagus* trees in Chile and we provide annotated genomes together with phenotypic characterization for 82 selected strains. We investigated the genetic structure and nucleotide diversity of this set of strains and re-analyzed other 23 previously published genomes. Overall, we provide evidence of a population structure in Patagonia that greatly expands previously known genetic structure in this region. Moreover, phenotypic clustering correlated well with genetic distances, where individuals from northern sites showed greater fermentation performance and high temperature tolerance than isolates from southern sites. The genomic data presented here broadens our knowledge of the genetics, ecology, and evolution of wild *Saccharomyces* strains.

## RESULTS

### S. eubayanus isolation and whole genome sequencing

In order to determine the presence and distribution of *S. eubayanus* isolates along the south western side of the Andes Mountains (which is within the Chilean territory), we sampled ten national parks and reserves between 2017 and 2018. The sites sampled correspond to primary forest spanning 2,090 km from Altos de Lircay National Park in central Chile (VII Maule Region, Chile) to Karukinka Natural Park in southern Chile (XII Magallanes Region, Chile) (**Figure 1A**). From these sites, we obtained 553 bark samples from *Nothofagus* and *Araucaria* trees, primordially *N. pumilio, N. antarctica, N. dombeyi* and *A. araucana.* Raffinose and ethanol media enrichment (Cubillos et al. 2019) allowed us to recover yeast colonies in 77% of the samples. Potential *Saccharomyces* strains were identified by sequencing the ITS and/or *GSY1* and *RPI1* RFLP (Peris et al. 2014; Peris et al. 2016). From these, 160 *S. eubayanus* strains were identified from different individual trees (**Table S1**, representing 28.9% of the samples), and in parallel, another set of 179 *S. uvarum* isolates were recovered (representing 37.9% of the samples), together with dozens of non-*Saccharomyces* species belonging to the *Lachancea, Kregervanrija, Kazachstania* and *Hanseniaspora* genera. Preliminary genotyping using *GSY1* RFLP analysis (Peris et al. 2014; Peris et al. 2016) suggested that all isolates belong to the PB lineage, except for isolate CL609.1 which showed a PA restriction pattern (data not shown). In general, we observed a pattern between yeasts and hosts, where *N. pumilio* and *N. antarctica* contained mostly *S. eubayanus* strains, while all but one isolate derived from *N. dombeyi* and *A. araucana* samples were identified as *S. uvarum* (**Table S1**). Moreover, the frequency of yeast isolates was higher towards southern regions (from Villarrica to Torres del Paine), where up to 90% of the bark samples yielded yeast colonies, and most of these belonged to the *Saccharomyces* genus (**Figure 1B**). On the contrary, a lower fraction of yeast colonies was obtained from Tierra del Fuego, likely due to the extreme environmental conditions found in this region. Overall, our results demonstrate the high frequency of the *S. eubayanus* species after latitude 33° in the western side of the Andes Mountains.

**Figure 1.**
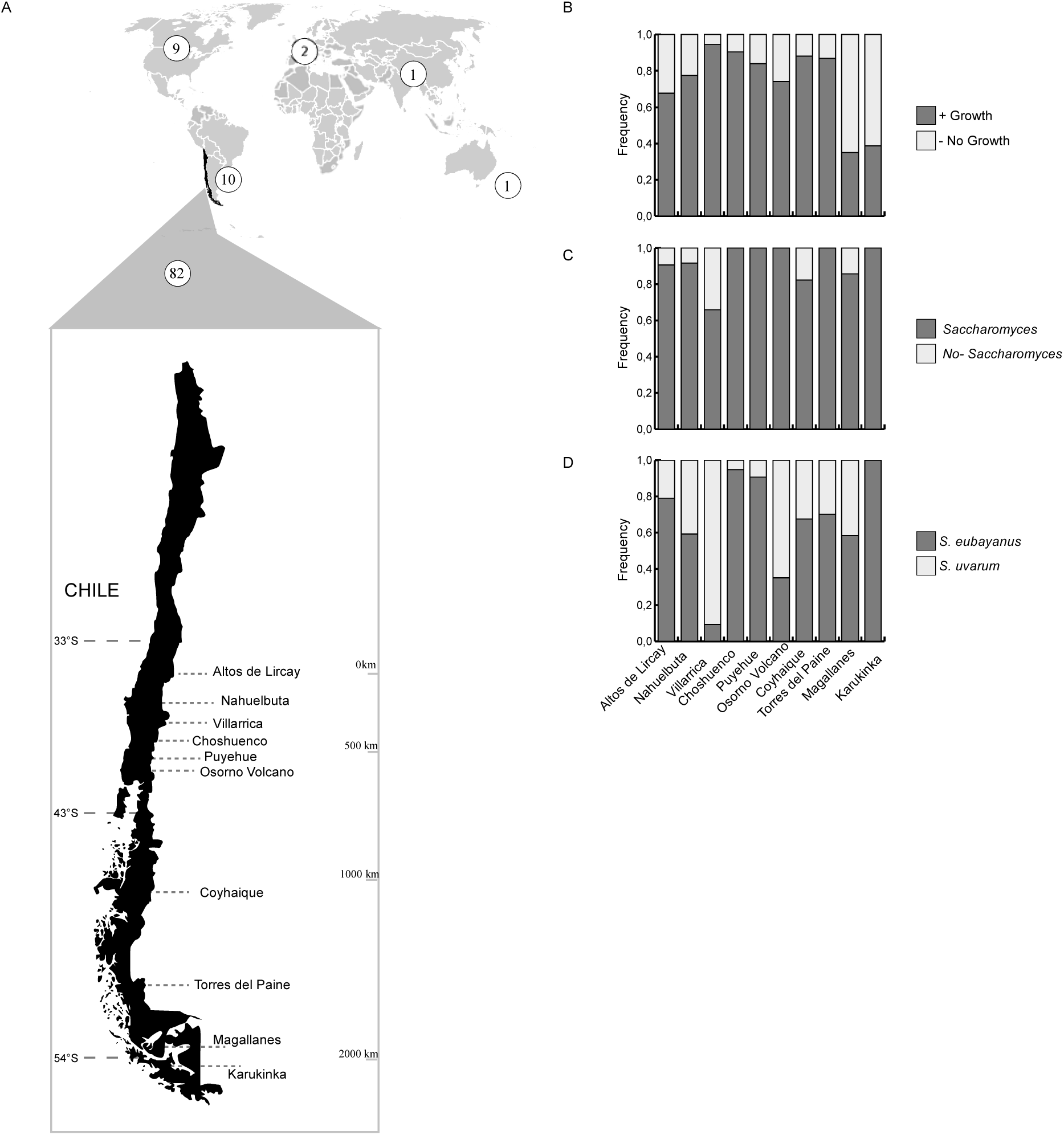
Geographic distribution and isolation frequency of *S. eubayanus* strains. (A) Map of the world depicting the number of available *S. eubayanus* sequenced genomes from around the world (white circles), together with the ten localities in Chile where the 82 strains sequenced in this study were isolated. Overall, a 2,090 km distance between sites was covered. Frequency of bark samples that yielded a (B) successful yeast isolation (dark grey) or no growth (light grey), a (C) *Saccharomyces* (dark grey) or other non-*Saccharomyces* genera (light grey), and a (D) *S. eubayanus* (dark grey) or *S. uvarum* (light grey) species.

To investigate the genomic variation and population structure of the *S. eubayanus* isolates, we sequenced the whole genomes of 82 strains, randomly selected, from the ten sampling sites using Illumina sequencing technology. Furthermore, to explore the genomic diversity across the entire geographic range of this species, we combined our dataset with all previously published genome from 23 strains obtained from the North America (9 strains), Argentina (10 strains) (Peris et al. 2016), China (1 strain) (Bing et al. 2014), New Zealand (1 strain) (Gayevskiy and Goddard 2016) and two lager genomes (Baker et al. 2015), maximising the geographical dispersal of the species. In parallel, FACS analysis revealed that all samples were diploids, except for CL609.1 and CL1005.1 that were found as haploid and tetraploid respectively (**Figure S1**). Indeed, all strains were able to sporulate, with the exception of CL609.1 (data not-shown).

On average, we obtained 25.9 million reads per sample, which were aligned against the CBS12357^T^ type strain reference genome (Brickwedde et al. 2018). From this, we obtained an average coverage of 164X, ranging from 17X to 251X (**Table S2**). A total of 229,272 single nucleotide polymorphisms, together with 19,982 insertions and deletions were found across the 82 *S. eubayanus* genomes collected in this study. The number of SNPs per strain ranged from 28,420 to 76,320 with strains CL619.1 and CL609.1 representing both extremes and which were isolated from Osorno Volcano and Puyehue neighbouring isolation sites, respectively (**Table S2a**). Overall, no strain isolated in this study represents a close relative to the type strain (obtained from Argentina), suggesting that the Andes Mountains represents a natural barrier between *S. eubayanus* populations (**Table S2b**). On average, across the 82 genomes we obtained 39,024 SNPs per strain relative to the reference genome, and a SNP was found on average every ∼300 bp. In parallel, we found on average 1,606 insertions and 1,677 deletions per isolate relative to the type strain. These results demonstrate that Chilean *S. eubayanus* populations are genetically distinct from those described in Argentina and world-wide.

### Population structure and admixed isolates

To examine whether the collected isolates comprise one panmictic population or various genetically distinct populations, we generated a neighbour joining phylogenetic tree based on 590,909 polymorphic sites (**Figure 2A**). The phylogeny obtained demonstrates that Chilean strains displayed different ancestry and mostly fell into three major clades: Patagonia B1 (PB-1), B2 (PB-2) & B3 (PB-3), each containing 31, 16, 25 strains, respectively. Nine strains fell outside the major PB lineage and might represent admixed strains (‘SoAm’). Also, one strain shares a recent common ancestor with the PA cluster suggesting that this strain might be a hybrid of the PB and PA lineages (**Figure 2B**). Each clade tends to contain strains obtained from neighbouring localities, suggesting a strong influence of geography on genetic differentiation. For example, strains obtained from southern (Coyhaique, Torres del Paine, Magallanes, Karukinka) and south-central Chile (Osorno Volcano) clustered in PB-1. Strains from central Chile (Altos de Lircay, Nahuelbuta and Villarrica) clustered in PB-2, and some strains from south-central Chile also clustered in PB-3 (Villarrica, Puyehue, Choshuenco, and Osorno Volcano). Interestingly, no isolates belonging to Patagonia A were found in Chile, and only a single isolate (CL609.1 from Puyehue) clustered near the PA branch, yet outside of this lineage.

**Figure 2.**
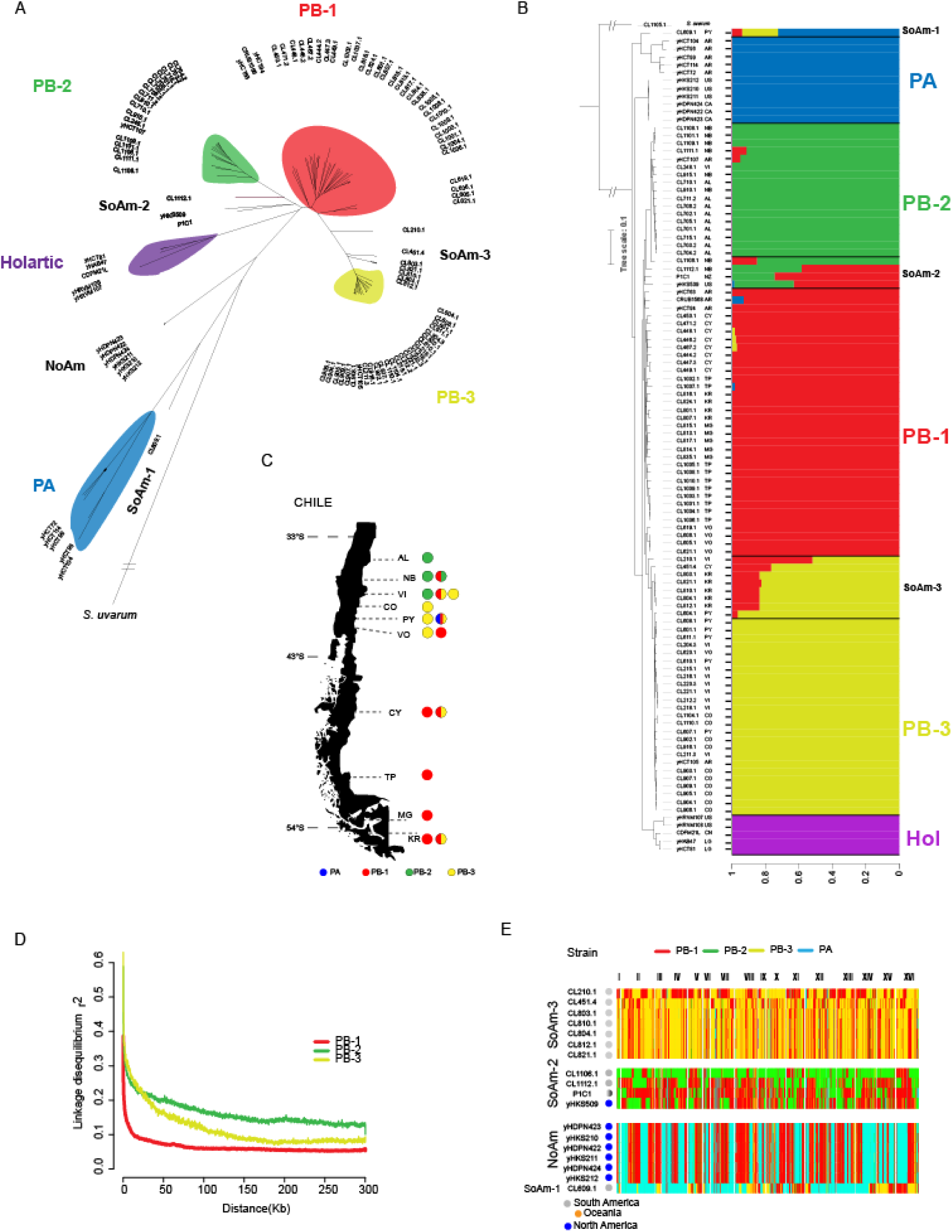
Phylogeny of *S. eubayanus*. (A) Whole genome Neighbour-joining tree built using 590,909 biallelic SNPs in 105 strains and manually rooted with *S. uvarum* as the outgroup. In all cases, bootstrap support values were 100% for all lineages. Three PB clades: PB-1 (red), PB-2 (green) and PB-3(yellow) and a single PA clade (blue) were identified, together with admixed strains between the different lineages. Branch lengths correspond to genetic distance. (B) Whole-genome Neighbour-joining tree of 105 strains built as in (A) together with the population structure generated with STRUCTURE. An optimum k = 5 groups was obtained. The geographic origin of each strain in depicted as follows: Canada (CA), United States (UN), China (CN), Lager (LG), AR (Argentina), New Zeland (NZ), AL (Altos de Lircay), NB (Nahuelbuta), Villarrica (VI), Choshuenco (CO), Puyehue (PY), Osorno Volcano (VO), Coyhaique (CY), Torres del Paine (TP), Magallanes (MG) and Karukinka (KR). Lineages distribution across sampling sites including PB lineages and SoAm admixed lineages. Linkage disequilibrium decay over distance (kb) expressed in terms of correlation coefficient, r2. LD decay for each window was estimated as the pairwise average for all SNPs pairs separated by no more than 100 kb. The PB-1 lineage shows the lowest LD values compared to any other population in our collection (E). Genome-wide ancestry for admixed strains. Bins of 100 SNPs were assigned in the admixed strains to the populations PB-1 (red), PB-2 (green), PB-3 (yellow) or PA (blue) based on sequence similarity.

To investigate the ancestry, we explored population structure using STRUCTURE. For this, we selected 9,885 SNPs evenly distributed across the whole genome. Among our analyzed strains, we identified three groups exhibiting different levels of mosaic or admixed genomes. This analysis indicated an optimum k = 5 groups (ΔK_5_ = 2,652, **Figure S2, Table S3**), highlighted by the presence of three main Patagonia B populations in Chile (**Figure 2B**).

Furthermore, sequence similarity using SNP data and principal component analysis (PCA) on the Patagonian populations validated the presence of three groups in addition to the Argentinian Patagonia A cluster (**Figure S3**). Most localities contained isolates belonging to one lineage and/or admixed groups, excepting for Villarrica and Osorno Volcano localities which harbour at least two lineages and/or admixed set of strains, representing sympatric geographic regions (**Figure 2C**). Overall, the phylogenomic, STRUCTURE and PCA analyses indicate that the *S. eubayanus* Patagonia-B clade found in Chile can be subdivided into three different lineages, PB-1, PB-2 & PB-3. Moreover, three groups of admixed strains (SoAm 1-3) were found, that together with the North American (NoAm), Holarctic and PA lineages shape the genetic structure of S. *eubayanus*.

In order to gain insight into the historical recombination events that affected the PB clade, we estimated linkage disequilibrium decay. Estimates of LD based on *r*^*2*^ values differed between the three PB lineages (**Figure 2D**). In particular, relatively low LD values were observed for all populations, but the maximum *r*^*2*^ and the rates of decay differed among populations. LD was detected over larger distances in the PB-3 and PB-2 populations, while LD decreased rapidly with increasing distance for the PB-1 population. Lineages showed a 50% LD decay of 2.9 kb, 29.1 kb and 22.5 kb in the PB-1, PB-2 and PB-3 populations, respectively, demonstrating a population-specific LD decay and greater recombination levels in the PB-1 population.

This could be explained by greater inbreeding or outbreeding rates in the clade. Indeed, *F*is values (Wright’s inbreeding coefficient) were significantly higher (*p*-value < 0.001, paired Student t-test) in PB-1 (average *F*is = 0.9482 CI = 0.9462 - 0.9503), compared to PB-2 and PB-3 (average *F*is = 0.9017 (CI = 0.8974 - 0.906 & *F*is = 0.8856 CI = 0.8795 - 0.8916, respectively), suggesting high inbreeding ratios in these populations (**Table S3b**). Moreover, the PB-1 level of recombination was similar to what is described in domesticated *S. cerevisiae* populations (Liti et al. 2009; Peter et al. 2018).

To determine how recombination events influenced the genomes of admixed strains and the level of genetic exchange between populations, we explored their mosaic genome compositions and genetic origins. Consequently, we generated similarity plots using 100 SNPs blocks and determined the closest genetic origin for each of the three groups of admixed strains from each population. The most interesting strain corresponds to CL609.1, which is a hybrid strain between the PA (58%), PB-1 (20%) and PB-3 (22%) lineages, where different segments fall into each lineage (**Figure 2E**). Admixed strains clustering nearby the PB-3 lineage represented another example where mosaic genomes between lineages were found. These strains could represent hybrids between PB-1 and PB-3, and most of these strains share the same regions from each population, and therefore suggesting a common ancestor (**Figure 2E**). Similarly, two admixed strains between PB-1 & PB-2 from Nahuelbuta showed different proportions of blocks of origins. While 79% of the genome of CL1106.1 is most similar to PB-2, 54% of the genome of CL1112.1 is most similar to genomes of individuals pertaining to PB-1. Similarly, the P1C1 strain from New Zealand corresponds to a hybrid originated from PB-1 & PB-2 lineages, while North American strains hybrids between PB-1 & PB-3, likely suggesting a migration out from Tierra del Fuego and the Patagonia. These results demonstrate the constant outcrossing and admixture between subpopulations and the success of the PB-1 branch by contributing to all of the admixed genomes analysed in our study.

### Highest nucleotide diversity in Tierra del Fuego populations

Characterizing patterns of genetic variation at the whole-genome level among populations can provide insights into signatures of selection (Hoban et al. 2016). Therefore, we calculated nucleotide diversity (π, which corresponds to the average number of nucleotide differences between individuals per site), genetic differentiation (*F*_*ST*_), and neutrality test statistics: Tajima’s D, Fu and Li’s D, and Fu’s F (**Table S3B**). Genome-wide nucleotide diversity (π) differed among PB populations (**Figure 3A**). PB-1 was more genetically diverse population (π= 0.00166750) than PB-2 (π= 0.000959206) and PB-3 (π= 0.000591578). These results are in agreement with PB-1 faster LD decay results and the broader geographic range where this clade is found. Interestingly, samples collected in Coyhaique showed the highest nucleotide diversity (π), in agreement with a high abundance of isolates in this locality (**Figure 3B)**. Indeed, southern localities belonging to PB-1 showed significantly greater π scores compared to the other two lineages, thus in our study PB-1 represents the most genetically diverse population (**Figure 3)**. On the other hand, PB-3 isolates from Choshuenco and Villarrica, located further north had the lowest levels of genetic diversity (**Figure 3B**). These results demonstrate a positive and significant (*p*-value < 0.05, Spearman correlation coefficient rs = 0.632) correlation between latitude and genetic diversity, i.e. the more southern the sampling the higher the genetic diversity found, and this is likely influenced by greater *N. pumilio* dispersal towards southern regions (**Figure 3C**).

**Figure 3.**
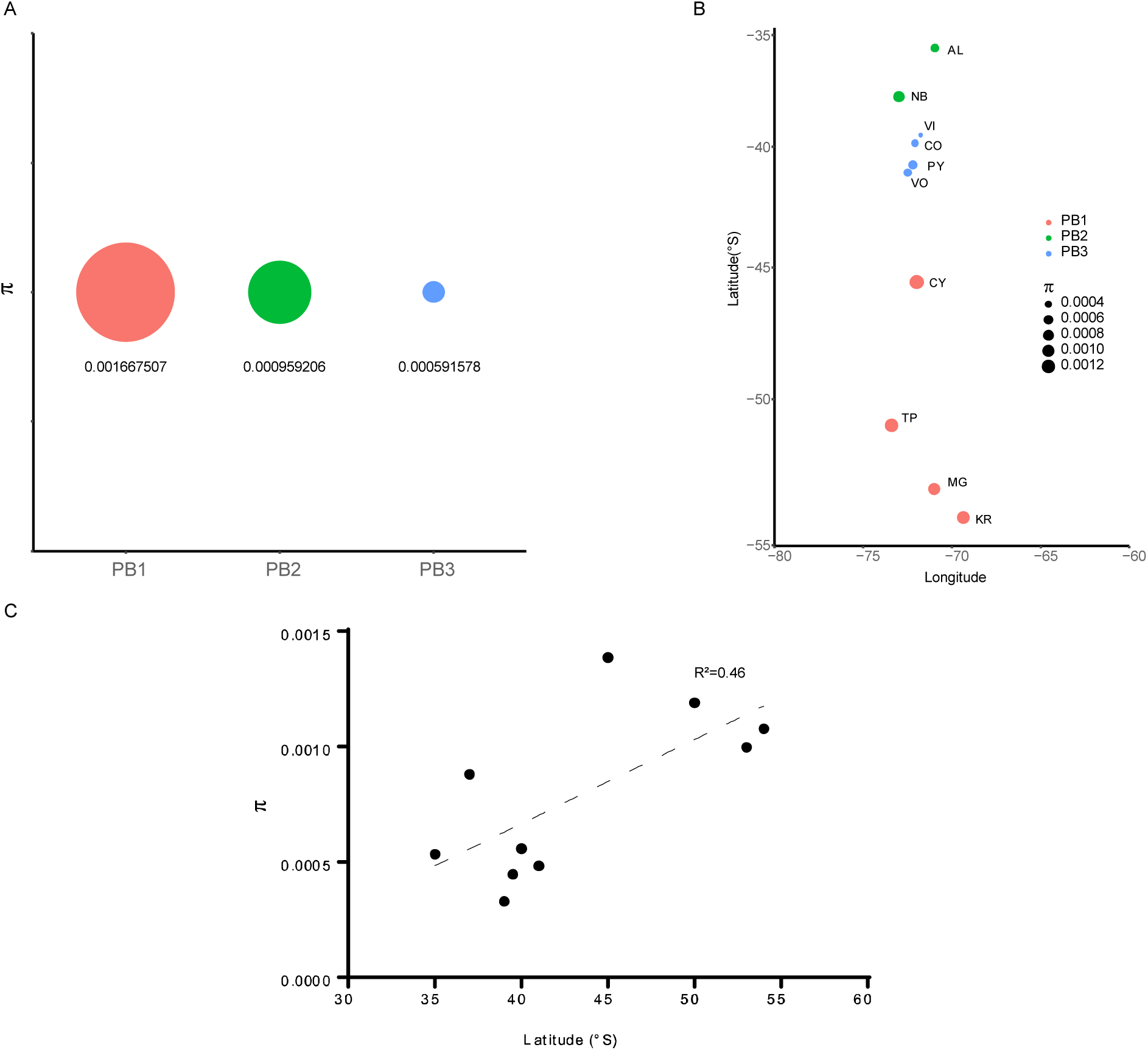
Nucleotide diversity in *S. eubayanus* across populations and localities in the western side of the Andes. Nucleotide diversity (π) in (A) PB populations obtained in this study and (B) localities across Chile. The geographic origin of each strain in depicted as follows: AL (Altos de Lircay), NB (Nahuelbuta), Villarrica (VI), Choshuenco (CO), Puyehue (PY), Osorno Volcano (VO), Coyhaique (CY), Torres del Paine (TP), Magallanes (MG) and Karukinka (KR). (C) Correlation between nucleotide diversity and latitude.

Subsequently, we estimated Tajima’s D values (measured as the deviation between pairwise differences and segregating sites). Tajima’s D’ scores differed between clades. Specifically, Tajima’s D for PB-2 & PB-3 were positive while this metric was nearly neutral for PB-1, suggesting balancing selection in the former case and neutral selection for the latest (**Table S3b**). To determine if the pattern of D’ scores was consistent across the whole genome, we plotted the Tajima’s D values along the genome for every 100 SNPs in non-overlapping windows. Interestingly, PB-3 showed a greater number of regions with positive Tajima’s D scores (>1) compared to the other two clades. This suggests that balancing selection may maintain allelic diversity in these regions (**Figure S4**).

The obtained *F*_*ST*_ values suggest that these populations are genetically different (*p*-value < 0.0001, **Figure S5, Table S4**). Our genome-wide analysis allowed us to find only a handful of regions sharing low Tajima’s D values between clades, yet all but one region in chromosome V exhibited high *F*_*ST*_ values. This genomic island found between PB-1 and PB-2 exhibited low Tajima’s D values and extremely low *F*_*ST*_ values (**Figure S4B & Figure S5**), suggesting a common genetic ancestry. This region contained four genes: *IRC22, MNN1, NOP16* and *PMI40*. From GO term analysis we found enrichment for ‘glycosilation’ process, due to the presence of *PMI40* and *MNN1*, the former encoding for an essential mannose-6-phosphate isomerase that catalyses the interconversion of fructose-6-P and mannose-6-P, while the latest encoding for an Alpha-1,3-mannosyltransferase. These results suggest that glycosilation could be under selection in these two populations.

We then assessed the degree of genetic differentiation and nucleotide diversity between the ten sampling sites per lineage. We found moderate to high significant *F*_*ST*_ values ranging from 0.16 to 0.88 (**Table S4**). Torres del Paine and Magallanes, two localities clustering in PB-1 and separated by ∼ 200 km had the lowest *F*_*ST*_ values, while Villarrica and Altos de Lircay (separated by 400 km) were the most genetically differentiated (**Table S4**). A mantel test showed a significant correlation between the geographic and the genetic distances (isolation by distance, *p*-value < 0.05) among sampling sites of the PB-1 lineage. The number of pairwise comparisons was insufficient to test for IBD for lineages PB-2 and PB-3. The positive IBD for PB-1 indicates limited effective dispersal within this lineage (**Figure S6**).

### Pangenome and genome content variation

In order to compare the genome content, we constructed the pangenome across all isolates. We identified 5,497 non redundant pangenomic ORFs in the species. Out of these, 5,233 ORFs are core systematically present in all of the isolates, while 264 are dispensable, being only found in subsets of strains. A PCA analysis of the presence/absence profile of these ORFs was used to visualize potential overlap between genome content similarities and SNPs distance **(Figure 4)**. In partial concordance with the phylogenetic tree, the branch harbouring the Chinese isolate, which also contained two North American isolates, represented the most divergent clade. The common grouping of Chinese and North American isolates together with their sequence similarity, suggests a recent migration event between Asia and America. Clear genetic differences were also found between PB-3 and the other clades. This suggests reduced admixture of these genomes, but a relatively recent separation. On the other hand, isolates from PB-1 and PB-2 distribute close to each other representing two halves of the same cluster, which can either suggest an extremely recent separation between the two clades, or continuous admixture. Interestingly, in our analysis PA isolates were extremely close to the PB-1 isolates, indicating extensive overlapping in genome content.

**Figure 4.**
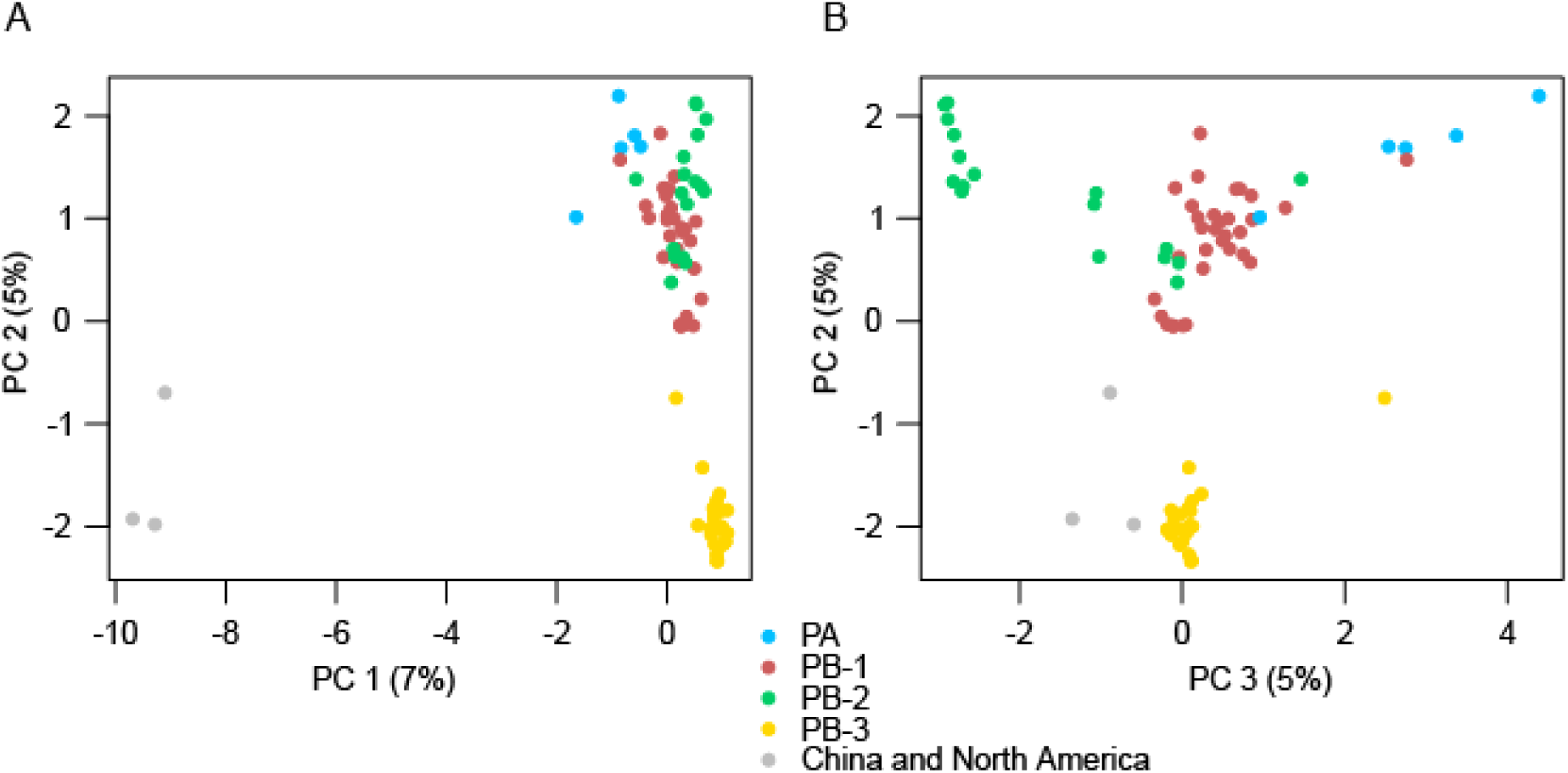
PCA of the gene’s presence-absence profiles. (A) Principal component analyses of the first three components show a reasonable level of concordance between sequence variation and genome content difference. Chinese/North American branch can be easily separated from the South American clades. Only non-mosaic 83 isolates from all populations were considered. (B) Principal component analyses considering only PB populations. Middle positioning of PB-1 mirrors the shape of the un-rooted phylogenetic tree based on the sequence divergence (Fig.2A). Interestingly, the PB-3 is the most separated clade, suggesting a lower level of outbreeding, while a partial overlap can be identified between the other clades.

To identify potential lateral gene transfer (LGT) events from other species, we compared the ORF sequences with an in-house database containing the ORFeome of 57 representative species

We identified nine ORFs present in a group of nine closely related isolates belonging to the PB-2 clade (CL248.1, CL701.1, CL702.1, CL705.1, CL710.1, CL711.1, CL715.1, CL910.1 and CL915.1), and six of the strains were isolated from the same locality and were assembled within a single contig for each isolate (*S. eubayanus* Region A). In two of these isolates (CL701.1 and CL248.1) these ORFs appear to be duplicated. SMART was used to identify known PFAM protein domains. These comprise putative proteins with an arginase domain, a MFS_1 (Major facilitator superfamily), a membrane transport protein domain, a Fungal specific transcription factor domain, a Gal4-like dimerization domain and a transmembrane one (**Figure S7**). In addition to these high-confidence hits, a search for homologies, also performed using SMART, indicated the presence of potential glucosidase domain in four of the other ORFs and a homing endonuclease on another one (**Figure S7**). The absence of matches from a search in the NCBI non-redundant database suggest the presence of orphan genes from an external un-sequenced donor species.

Furthermore, we also identified 64 private ORFs in the Chinese/North American clade. Out of 64 such ORFs, at least 20 correspond to orthologs of other genes found in *S. eubayanus* (showing an identity percentage between 94% and 76%). For the majority of such ORFs no potential donor species has been found. That being said, one of these ORFs was identified as a *MALS* gene which encodes for a maltase enzyme also found in *S. uvarum* x *S. eubayanus* hybrids (*S. bayanus*). These ORFs are not found clustered together, but rather scattered across several contigs in the assemblies. A phylogenetic analyses performed gene by gene indicates the potential donor species to have an intermediate distance between *S. eubayanus* and *S. arboricola*, although the presence of more than one donor cannot be categorically excluded. It is possible that different subset of these ORFs have different evolutionary origins. Among these ORFs, we could also find ancestral segregating genes, paralogs of other *S. eubayanus* genes and lost in the lineage from which the South American clades originated.

### Phenotypic diversity among S. eubayanus isolates

*S. eubayanus* phenotypic diversity was assessed in a set of 88 isolates, representative of the different clades found across the American continent. High-resolution growth quantification was conducted under eight environmental conditions. These conditions included different growth temperatures, salt resistance, glucose and maltose utilization and antimicrobial compounds. Three growth variables were extracted: the lag phase, growth rate (μmax) and maximum OD (maxOD) (**Table S7a)** generating phenotypic data for 24 attributes from the growth curves. We found a significant correlation between μmax and maxOD, and therefore we focused on these two as fitness parameters and compared them across isolates & conditions. We found that bivariate correlations among traits vary considerably (**Table S7b**). For example, the conditions NaCl, and CuSO_4_ showed a positive μmax correlation (Pearson r = 0.47, p-value < 6 × 10^−9^), where both traits likely share similar molecular pathways (Dhar et al. 2013). Conversely, there was no significant μmax correlation between NaCl and Hygromycin resistance (Pearson r = 0.12, *p*-value = 0.14). NaCl and 34°C represented the two phenotypes exhibiting the greatest phenotypic coefficients of variation (47% and 62%) for μmax and maxOD, respectively, suggesting ecological niche differentiation and mutation accumulation in these polygenic traits across the strains.

We then generated a clustered heat map of the phenotypic correlations between yeast isolates across all traits for μmax (**Figure 5**). Phenotypic clustering was mostly driven by geography, rather than by genetic relationship. North American Holarctic strains clustered separately and showed low μmax and maxOD scores across traits. The CBS12357^T^ strain clustered together with PB-1 strains from Torres del Paine. This clustering was further supported by the principal component analysis, both of which produced the main groups split by localities, rather than lineages, where only PB-2 isolates grouped together (**Figure S8**). Specifically, only for certain phenotypes we found differences among localities (**Table S8**).

**Figure 5.**
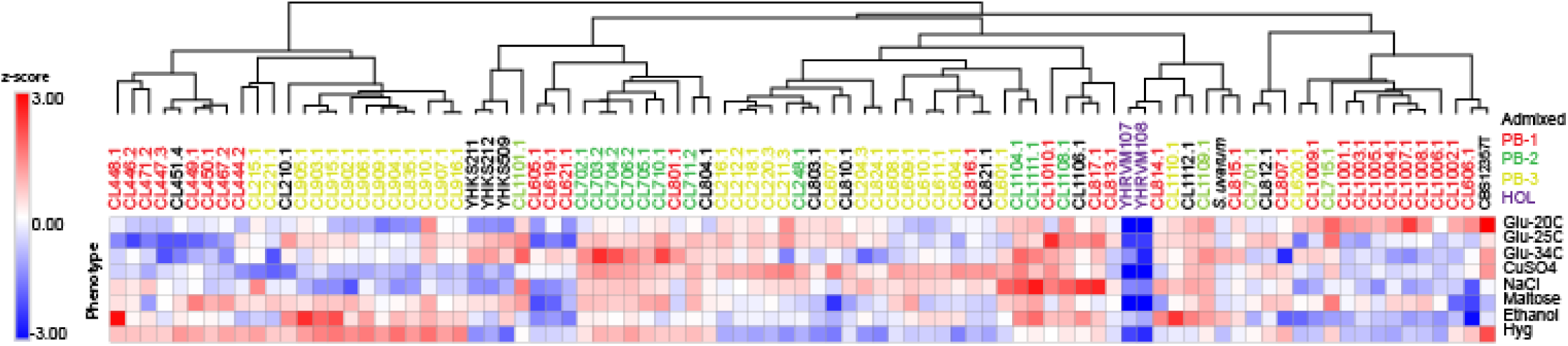
Phenotypic diversity in *S. eubayanus*. Heat map depicting the phenotypic diversity in *S. eubayanus* obtained from eight different conditions assessed in microcultures. Strains are grouped by hierarchical clustering and names & colours indicate the clade. The heat maps were elaborated based on z-scores within each phenotype.

For example, northern isolates obtained from Altos de Lircay and Nahuelbuta had greater μmax when grown at high temperatures (34°C) than isolates from other localities (*p*-value < 0.05, Tukey test). This suggests that northern isolates might be adapted to warmer climates. Altogether, our results demonstrate a relatively low phenotypic diversity across traits and strains, and most of this diversity can be explained by habitat adaptation rather than by phylogenetic history.

### Isolates from lower latitudes exhibit greater fermentation performance

Given the importance of *S. eubayanus* in lager brewing, we then evaluated the fermentation performance of the same set of strains previously phenotyped and used the W34/70 lager strain as fermentation positive control. The conditions included 12°P wort at 12°C in 50 mL (micro-fermentations batches). For this, CO_2_ loss was recorded every day and metabolite consumption (glucose, fructose, maltose and maltotriose) and production (glycerol and ethanol) was estimated at the end of the fermentation process. In all cases, the lager control showed better fermentation performance compared to the majority of *S. eubayanus* isolates (*p-*value < 0.05, t-test, **Table S9a**), except for five strains from Altos de Lircay and Villarrica. This strains showed not-significantly differences in CO_2_ lost levels compared to the lager control (*p*-value > 0.05, Student test), demonstrating the greater beer fermentation potential of these strains. Overall, we observed that isolates obtained at lower latitudes (Central region) lost significantly greater CO_2_ levels than individuals obtained at higher latitudes (extreme South). For example, Magallanes isolates belonging to PB-1 showed 2X lower fermentation performance than isolates obtained from Altos de Lircay (PB-2), which are separated by approximately 1,800 km (**Table S9b**). Indeed, we found a significant correlation (Pearson r = 0.566, *p*-value < 0.001) when we compared latitude vs CO_2_ lost (**Figure 6A**) and significant differences between localities (**Table S9b**). Furthermore, we found that fermentation performance was directly correlated with maltose sugar consumption (Pearson r = 0.52, *p*-value < 6 × 10^−8^, **Table S10, Figure 6B**), and ethanol production (Pearson r = 0.56, *p*-value < 2 × 10^−6^).

**Figure 6.**
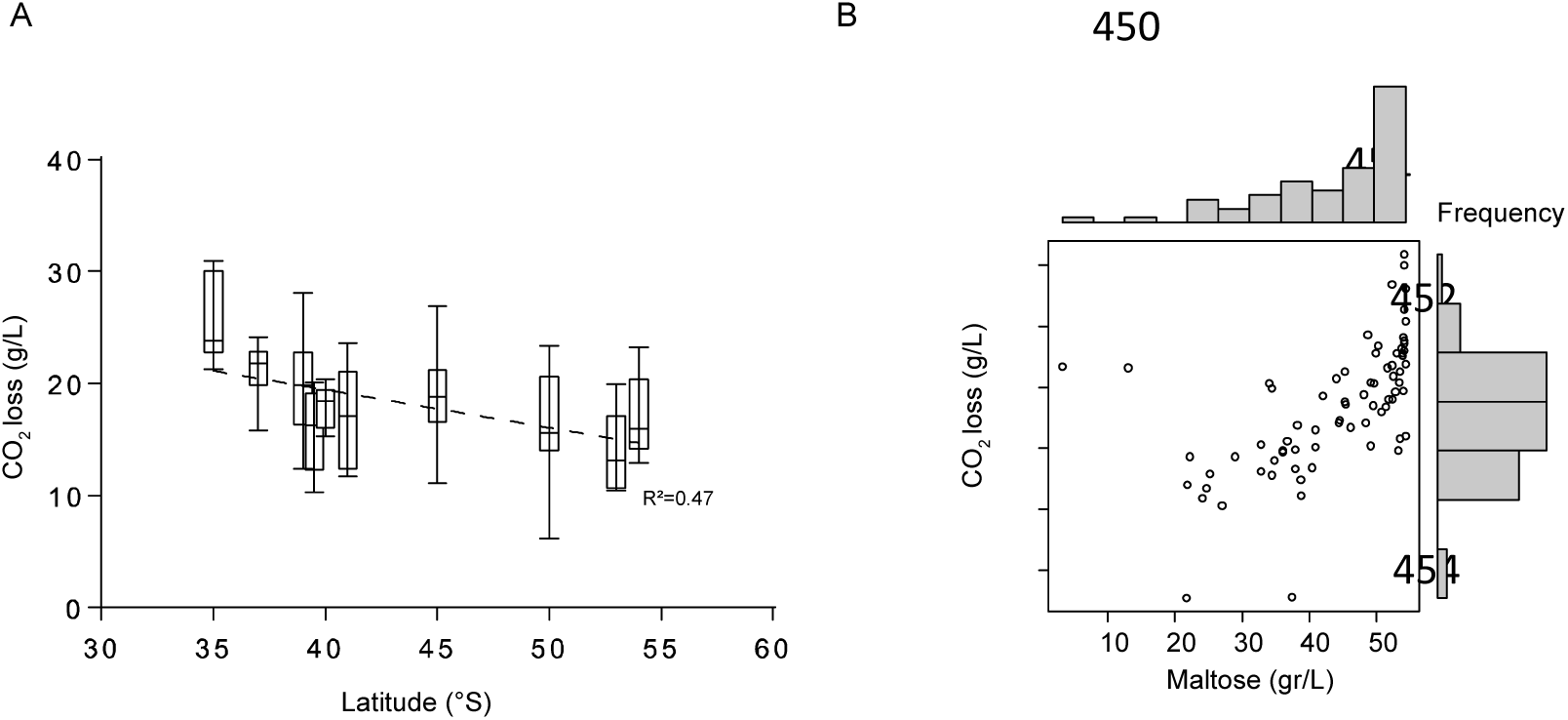
Fermentative profile of *S. eubayanus* strains. (A) CO2 loss levels represent the fermentative capacity of wild isolates obtained from microfermentations. Error bars denote the standard deviation (B) Maltose consumption was directly correlated with CO2 loss and latitude of the origin of the isolate.

In order to determine the molecular basis of differences in maltose consumption and fermentation performance, we evaluated the impact of polymorphisms and indels within the *AGT1* gene. This gene encodes for a high-affinity maltose and maltrotriose transporter in *S. pastorianus* and is responsible for the maltotriose consumption under fermentative conditions (Nakao et al. 2009). Three copies along the reference genome were found in chromosomes V (DI49_1597), XIII (DI49_3958) and XVI (DI49_5193), exhibiting a sequence identity above 55% and hereafter denominated as *AGT1_V, AGT1_XIII* and *AGT1_XVI*. In fact, analysing sequence variation across strains revealed that phylogenetic trees differed between *AGT1* copies (**Figure S9A**). Interestingly, we found that some strains do not carry all three functional copies of the Agt1 protein. This loss of function is caused by deletions that generate truncated proteins. For the *AGT1_V* encoded transporter, truncation occurs after 50 aminoacids, for the *AGT1_XIII* encoded transporter it occurs after 180 aminoacids, and for the *AGT1_XVI* encoded transporter it occurs after 293 aminoacids (**Figure S9B**). Interestingly, PB-2 isolates did not carry any of the truncations, however many non-synonymous polymorphisms were found within the *AGT1* sequences. Indeed, docking simulations indicate reduced maltose and maltrotriose binding capacity in the truncated proteins in comparison to the non-truncated Agt1p copies (**Figure S9C**, *p*-value < 0.001 Student-test). Nevertheless, we did not find a direct correlation between fermentation performance and a specific truncated Agt1p copy or a significant correlation between fermentation performance and number of copies of the functional Agt1p transporter. These results suggest that other genomic regions might be involved in maltose uptake generating the observed differences in fermentation performance.

## DISCUSSION

Our results, as well as others (see (Libkind et al. 2011; Peris et al. 2014) strongly suggest that *S. eubayanus* is preferentially found in association with *Nothofagus* trees, particularly *N. pumilio* (the most cold-adapted *Nothofagus* (de Porras et al. 2012; Hinojosa et al. 2016b). Under this scenario, the biogeographic history of *S. eubayanus* should be strongly correlated with the *Nothofagus* dispersal history across the globe. Interestingly, the isolation frequency of *S. eubayanus* was correlated with the latitude. PB-1 is located at higher latitudes (and lower altitudes) and the isolation frequency of this lineage was lower than that of the other two. Overall, PB-2 and PB-3 were easily recovered from the environment and were specifically associated with high altitude *N. pumilio* trees. These results are in agreement with those reported for *S. eubayanus* in Argentina (Eizaguirre et al. 2018). The lower frequency of isolates found in samples from Tierra del Fuego could be due to the extreme environmental conditions confronted in this part of the continent, with average temperatures below 5°C throughout most of the year (Ponce and Fernández 2014). However, the extended distribution and higher abundance of *N. pumilio* in Tierra del Fuego compared to northern areas may facilitate *S. eubayanus* survival, range distribution, and habitat colonization (Hildebrand-Vogel et al. 1990) increasing population size and genetic diversity. The genus *Nothofagus* was originated during the late Cretaceous-Early Tertiary interchange (ca. 135 MYA); between Southeast Asia and Australia, from a “fagalean” complex in Southeast Asia (Hill 1992b; Hinojosa et al. 2016b). At the time, Antarctica was in a northern position connecting South America, Tasmania, Australia and New Zealand; and had a warm-humid climate (=”mesothermal conditions”, sensu (Hinojosa et al. 2016b)). Then, a step-wise dispersion-colonization wave following a westward direction reached South America. This scenario is consistent with a dispersal of the *S. eubayanus* and *S. uvarum* ancestor across the South Hemisphere (Hill 1992a; Hershkovitz 1999; Dutra and Batten 2000).

The phylogenetic analysis presented here of wild *S. eubayanus* demonstrates that the Patagonia B lineage is actually composed of three clean lineages: PB-1, PB-2 and PB-3. Interestingly, no strains belonging to the Patagonia A lineage were found amongst our samples despite this lineage being highly abundant in Argentinian Patagonia (Eizaguirre et al. 2018; Langdon et al. 2019). From the linkage disequilibrium analysis we found smaller linkage blocks and higher *F*is values in the PB-1 cluster, compared to PB-2 and PB-3. This suggests that meiosis and inbreeding are more frequent in PB-1. Additionally, the nucleotide diversity of PB-1 was higher than that of the other populations, and this was supported by greater genetic diversity among individuals sampled from Coyhaique, Torres del Paine, Magallanes and Karukinka locations. Individuals from these sites had high nucleotide diversity which is interesting given that dispersal of *N. pumilio* is also greater than in northern regions. Nevertheless, southern regions only contained individuals from a single lineage, contrasting northern sites which harbour individuals from more than a population. Interestingly, a subgroup of the PB-1 isolates is found at lower latitudes, which means that PB-1, PB-2 and PB-3 are sympatric. Indeed, our STRUCTURE and pan-genome analyses provide evidence of contact and admixture between PB-1 and the other two. That being said, some of these admixed strains have migrated to other regions around the world, including North America, China and Oceania. Overall, all of the admixed strains from Chile, New Zealand and North America contained regions from the PB-1 lineage, suggesting an ‘out-of-Patagonia’ origin for most strains in the Holarctic lineage, and even Oceania, and demonstrating the success of the PB-1 lineage.

The genetic diversity reported here for the PB South American clades is higher than that reported for *S. eubayanus* in the Northern Hemisphere (Peris et al. 2016). Indeed, our results resemble those obtained in the sister species for *S. uvarum*, where similar sequence divergence among *S. uvarum* populations were described (Almeida et al. 2014). Interestingly, based on the levels of genetic diversity and heterozigosity, our results support the idea that PB-1 *S. eubayanus* from Tierra del Fuego is the oldest population in Patagonia, and likely within the species. In fact, this lineage shows high levels of hybridization/introgression into northern populations (see Fig 2). This scenario is also consistent with the idea that *S. eubayanus* cryotolerance evolved recently, as *N. pumilio* (its preferred environment) became secondarily adapted to cold during the orogenesis of the Andes, a relatively young mountain range (de Porras et al. 2012; Hinojosa et al. 2016a; Horton 2018). Also, it is now accepted that the distribution of *Nothofagus* (and subsequently

*S. eubayanus*) was already established in Patagonia before the onset of the last glacial maximum (ca 20,000 years ago), whose ice sheets covered most land masses south of 41°. Thus, there was massive floristic recolonization and ecological succession after this period (Sersic et al. 2011; de Porras et al. 2012). Our results are consistent with this second scenario (colonization from peripheral glacial refugia from the South) since we found lower genetic diversity in populations located in central Chile (Altos de Lircay & Nahuelbuta) than in populations found in southern Chile (Coyhaique, Torres del Paine, Magallanes and Karukinka). Interestingly, a different pattern would be observed on the other side of the Andes (Argentinian territory), where a lower number of peripheral glacial refugia occurred and most of the diversity would originate from Valleys refugia in northen sites (Sersic et al. 2011). Indeed, in Argentina a greater *S. eubayanus* genetic diversity is reported north of 41°, where different lineages congregate in a single geographic location (Langdon et al. 2019). Thus, different glaciation refugia would have shaped the current genetic diversity and dispersal of *S. eubayanus* populations in Patagonia.

Our phenotypic assay demonstrates a relatively low phenotypic diversity between populations and localities, providing evidence that these populations might have experienced mild selective pressures in the environments here evaluated. Individuals located in northern populations grew much faster at higher temperatures than individuals from the south, in agreement with the idea of local adaptation. High temperature growth has been extensively studied in *S. cerevisiae* isolates (Steinmetz et al. 2002; Parts et al. 2011; Yang et al. 2013; Wilkening et al. 2014), where several natural variants were mapped down to the gene level. The fact that fermentation capacity was negatively correlated with latitude represents a striking to the low phenotypic diversity found under microcultivation conditions, suggesting that isolates found in northern regions could represent potential new strains for the brewing industry. Specifically, strains found at lower latitudes showed greater CO_2_ release and maltose consumption, contrasting with fermentation performances obtained in the east side of the Andes where northern isolates belonging to the PA lineage showed lower fermentation performances (Eizaguirre et al. 2018). Selection in maltose transporter could impact sugar assimilation (Brickwedde et al. 2018). In this context, the *AGT1* gene encodes for a high-affinity maltose and maltrotriose transporter (Alves et al. 2008) and is responsible for this capacity in *S. pastorianus. S. eubayanus* strains carry different putative copies of *AGT1*, however none of these are directly correlated with fermentation performance nor with the transport of maltotriose. Yet, overexpression of a Holarctic allelic variant of se*AGT1* was able to confer maltotriose consumption (Baker and Hittinger 2019). As expected, in our study no wild *S. eubayanus* isolate was found to consume maltotriose; this was not surprising given that *S. cerevisiae* is the only *Saccharomyces* species in the genus able to utilize maltrotriose as its carbon source (Krogerus et al. 2015 2018).

In sum, our results provide evidence of an ‘Out-of-Patagonia’ dispersal in the *S. eubayanus* species and that this dispersal is responsible for the current extensive genetic diversity found in the species. The majority of the *S. eubayanus* strains collected around the world belong to the Patagonian cluster, even a subset of those recently found in China, supporting a successful colonization from Out-of-Patagonia towards the Northern Hemisphere and Oceania. Finally, our data in *S. eubayanus* together with previous evidence in *S. uvarum* (Almeida et al. 2014) could possibly suggest that the ancestor of both species originated in the South Hemisphere, rather than China. However, the current available data is insufficient to draw further conclusions regarding the evolutionary history of the two species and future studies and evidences are needed to support these results.

## MATERIALS AND METHODS

### Sample areas and yeast isolation

Bark samples from ‘lenga’ (*Nothofagus pumilio*), coigüe (*N.dombeyi*) and ‘ñirre’ (*N. Antarctica*) and *Araucaria araucana* were obtained aseptically from ten sampling sites in Chile (collection date, **Figure 1**): National Park Altos de Lircay (January 2018, 35°36’34”S, 70°57’58”W), Nahuelbuta National Park (February 2018, 37°47’33”S, 72°59’53”W), Villarrica National Park (January 2017, 39°28’52”S, 71°45’50”W), Choshuenco National Park (January 2018, 39°50’2”S, 72°4’57”W), Antillanca National Park (November 2017, 40°46’23”S, 72°12’15”W), Vicente Pérez Rosales National Park (November 2017, 41°6’15”S, 72°29’45”W), Coyhaique National Reserve (February 2017, 45°31’23”S, 71°59’19”W), Torres del Paine National Park (February 2018, 50°56’32”S, 73°24’24”W), Magallanes National Reserve (January 2018, 53°8’45”S, 71°0’12”W) and Karukinka Natural Park (January 2018, 54°6’4”S, 69°21’24”W). All sampling sites were located at least five km from human settlements.

For each site, at least 25 bark samples of about 1g and 20 × 1 mm were obtained and immediately incubated in a 15 mL tube containing 10 mL of enrichment media. The media contained 2% yeast nitrogen base, 1% raffinose, 2% peptone and 8% ethanol (Sampaio and Goncalves 2008). Overall, 553 samples were collected (**Table S1**). Samples were incubated for two weeks at 20°C without agitation and were subsequently vortexed and plated (5 μL) onto YPD agar (1% yeast extract, 2% peptone, 2% glucose and 2% agar). Isolated colonies were stored in glycerol 20% v/v and stored at −80°C in the Molecular Genetics Laboratory yeast collection at Universidad de Santiago de Chile.

### *Saccharomyces eubayanus* identification *and FACS analysis*

We amplified and sequenced the internal transcribed spacer region (ITS) to identify colonies to the genus level. For this, ITS1 and ITS4 primers (J White et al. 1990) were used and we classified as *Saccharomyces* fragment sizes ranging between 830 and 880 bp (Pham et al. 2011). Species identification was conducted using the polymorphic marker *GSY1* and *RIP1* through amplification and enzyme restriction (see details in (Peris et al. 2014)). Then, restriction fragment length polymorphism were performed using the restriction enzymes *HaeIII* and *EcoRI* as previously described (Peris et al. 2014). Colonies were classified based on restriction patterns as either *S. eubayanus, S. uvarum* or *S. cerevisiae* (Peris et al. 2014). In many cases, species identification was confirmed by Sanger-sequencing of the ITS region, which was attained using a BLASTN against the Genbank database under 100% identity as threshold.

DNA content was analysed using a propidium iodide (PI) staining assay. Cells were first pulled out from glycerol stocks on YPD solid media and incubated overnight at 30 °C. The following day a small portion of each patch was taken with a pipette tip and transferred in liquid YPD in a 96-well plate and incubated overnight at 30 °C. Then, 3 μl were taken and resuspended in 100 μl of cold 70% ethanol. Cells were fixed overnight at 4 °C, washed twice with PBS, resuspended in 100 μl of staining solution (15 μM PI, 100 μg/ml RNase A, 0.1% v/v Triton-X, in PBS) and finally incubated for 3 h at 37 °C in the dark. Ten thousand cells for each sample were analysed on a FACS-Calibur flow cytometer using the HTS module for processing 96-well plates. Cells were excited at 488 nM and fluorescence was collected with a FL2-A filter. The data collected were analysed in R with flowCore (Hahne et al. 2009) and flowViz (Sarkar et al. 2008) and plotted with ggplot. The highest density value of FL2-A was associated with the ploidy level of G1 cells, thus cells that are not dividing, and used for inferring the ploidy state of the sample. FL2-A values between 60 and 110 for G1 cells were associated with haploid state, FL2-A values between 120 and 220 were associated with diploid state and FL2-A values between 290 and 400 were associated with a tetraploid state.

### Sequencing, Reads processing and Mapping

DNA was obtained using a Qiagen Genomic-tip 20/G kit (Qiagen, Hilden, Germany). The library prep reaction used was a 100x miniaturized version of the Illumina Nextera method. In this prep, 1.6 ng of total DNA mass is tagmented in a 5X diluted Tagmentation reaction. The 0.5 μL reaction was quenched by 0.5% SDS(0.125% final concentration) at room temperature for 5 minutes. After quenching, 125 nL of a P5 sequencing barcode and 125 nL of a P7 sequencing barcode were added to the 0.625 nL reaction. In order to amplify the library of inserts, 24.125 μL of 1X KAPA Library Amplification Master Mix were added to the reaction. The library went through 15 cycles of PCR to add the barcodes to then amplify the library to a concentration >4 nM. The libraries were then normalized and pooled according to the Illumina standard operating procedure and sequenced on a NextSeq 500/550 High Output Kit v2.5 (300 Cycles) flow cell.

Read quality and summary statistics were examined using FastQC 0.11.8 (Andrews 2014). Reads were processed with fastp 0.19.4 (low quality 3’ end trimming, 37 bp minimum read size) (Brickwedde et al. 2018; Chen et al. 2018). We also obtained publicly available sequencing reads of *S. eubayanus* (Bing et al. 2014; Gayevskiy and Goddard 2016; Peris et al. 2016; Brickwedde et al. 2018) and *S. pastorianus* (Baker et al, 2015} from the SRA database, which were processed similarly, i.e. visual inspection with FastQC and processing adaptors, low quality 3’ ends, and read size, with fastp. Processed reads were aligned against the *Saccharomyces eubayanus* CBS12357^T^ reference genome (Brickwedde et al. 2018) using BWA-mem (options: -M -R)(Li 2013). Mapping quality and overall statistics were collected and examined with Qualimap (García-Alcalde et al. 2012). Summary statistics are shown in **Table S2**. Sorting and indexing of ouput bam files were performed using SAMTOOLS 1.9 (Li et al. 2009). A *S. uvarum* isolate (CL1105) isolated from Nahuelbuta was also mapped to the *S. eubayanus* and *S. uvarum* CBS7001 genome (Scannell et al. 2011; Almeida et al. 2014) for phylogenetic analysis.

### Variant calling

Mapping files were tagged for duplicates using MarkDuplicates of Picard tools 2.18.14 (http://broadinstitute.github.io/picard/). Variant calling and filtering was done with GATK version 4.0.10.1 (DePristo et al. 2011). More specifically, variants were called per sample and chromosome using HaplotypeCaller (default settings), after which variant databases were build using GenomicsDBImport. Genotypes for each chromosome were called using GenotypeGVCFs (-G StandardAnnotation). Variant files were merged into one genome-wide file using MergeVcfs. This file was divided by SNP calls and INDEL calls using SelectVariants. We applied GATK recommended filters to both variant files, i.e. for SNPs “QD < 2.0 ‖ FS > 60.0 ‖ MQ < 40.0 ‖ MQRankSum < −12.5 ‖ ReadPosRankSum < −8.0”, and for INDELS “QD < 2.0 ‖ FS > 200.0 ‖ ReadPosRankSum < −20.0”. We then processed the SNPs VCF file with vcftools (---minQ 30, --max-missing 1, --max-alleles 2 (Van der Auwera et al. 2013). Furthermore, we applied a stricter criteria to filter heterozygous calls using bcftools view (-e ‘GT=”0/1” & QUAL<7000 & AC=1’) version 1.9 (Li et al. 2009)}. In addition, the effect of each variant was assessed and annotated with SnpEff version 4.3t (Cingolani et al. 2012), using an updated version of *S. eubayanus* gene annotations (Brickwedde et al. 2018)

### Phylogeny analyses

We obtained a phylogenetic tree using 590,909 biallleic SNPs. VCF files were imported to R (version 3.5.2)(Development Core Team) and converted to genlight objects with vcfR version 1.8.0 (Knaus and Grunwald 2017). A bitwise distance matrix was calculated with the package poppr version 2.8.1 (Kamvar et al. 2014), and a neighbour-joining tree was built using the function aboot, using 1000 bootstraps. Trees were visualized in the iTOL website (http://itol.embl.de). A thinned version of the VCF file was generated with vcftools 0.1.15 (--thin 1000)(Danecek et al. 2011), containing 9,885 similarly-spaced SNPs. Structure was run on this dataset five times for K values ranging from 3 to 7, with 10,000 burn-in and 100,000 replications for each run and using admixture model, infer alpha, lambda =1, fpriormean =1, unifprioralpha 1, alpha max 10. The structure-selector website was used to obtain the optimal K values (http://lmme.qdio.ac.cn/StructureSelector/) (Li and Liu 2018) according to the Evanno method (Evanno et al. 2005) and to obtain the final results for each K, which were plotted using CLUMPAK (Kopelman et al.). The resulting diagrams were visualised using structure plot (http://omicsspeaks.com/strplot2/)(Ramasamy et al. 2014). In addition, we performed clustering analyses of the same samples by using Discriminant Analysis of Principal Components (DAPC) of the adegenet R package version 2.1.1 (Jombart 2008) run with PB-1, PB-2 and PB-3 as population priors. Linkage disequilibrium (LD) between lineages was calculated using variants belonging to each Patagonia B population. LD decay was estimated by calculating R2 values using with vcftools (----geno-r2 --ld-window-bp 100000), which were imported into R to calculate a regression according to (Hill and Weir 1988), for which the half decay was estimated (Ldmax/2).

### Population Genetics

We estimated standard population parameters π, Tajima’s D, Fu and Li’s D, and Fu’s F using the R packages PopGenome 2.6.0 (Pfeifer et al. 2014). Values of *F*_st_ were estimated with StAMPP 1.5.1 (Pembleton et al. 2013) to obtain 95% confidence intervals by performing 5,000 bootstraps. Populations were designated as PB-1, PB-2, PB-3, or by sampling into each geographical location from the same lineage (for example, only PB-1s were compared between localities).

The variants of the mosaic *S. eubayanus* strains were split to bins of 100 SNPs (on average ∼5kb windows) and each bin was assigned to either of the populations (i.e. PB-1, PB-2, PB-3, or PA) using adegenet’s hyb.pred function. This algorithm uses DAPC to estimate membership probability of a hybrid dataset to a known cluster (populations). The Argentinian strains yHCT104 and yHCT72 were used as PA representative members.

The R package hierfstat (Goudet 2005) was used to calculate *F*_is_, Hs, and Ho by using the basic.stats function. Bootstrapping per loci on each population’s *F*_is_ was done using hierfstat’s boot.ppfis, obtaining the 50th and 97.5th quantiles after 50000 boostraps. To perform a Mantel test, first the Nei’s genetic distances between subpopulations (considering localities) was calculated with the R package StAMPP (Pembleton et al. 2013). Euclidean distance between localities was calculated using latitude and distances coordinates with R ‘dist’ function. Randel test was performed using the ade4 R package (Dray and Dufour 2007).

### Pangenome

Isolates were assembled with assembled with Spades using k from 21 to 67. To detect the non-reference material we used the custom pipeline based on the method described in (Peter et al. 2018). LRSDAY software (Yue and Liti) was used to annotate the non-reference material. The newly annotated ORFs have been added to the reference ORFs and a custom pipeline, also based on methods from (Peter et al.) was used to collapse ORFs with identity percentage over 95, selecting an unique reference for each groups of allelic variants to obtain a list of non-redundant pangenomic ORF sequences. Confirmation of presence of these ORFs has been obtained by mapping the reads of each strain to the set of pangenomic ORFs using BWA mem with the option – U 0. Filtering was performed with samtools with options –bSq 20 –F260.

### Strains Phenotyping and Fermentations

The microcultivation phenotyping assay of the *S. eubayanus* strains was performed as previously described (Kessi-Perez et al. 2016). Briefly, isolates were pre-cultivated in 200 μL of YNB medium supplemented with glucose 2% for 48h at 25°C. For the experimental assay, strains were inoculated to an optical density (OD) of 0.03–0.1 (wavelenght 630 nm) in 200 uL of media and incubated without agitation at 25°C for 24 h (YNB control) and 48 h for other conditions in a Tecan Sunrise absorbance microplate reader. OD was measured every 20 minutes using a 630 nm filter. Each experiment was performed in quadruplicate. Maximum growth rate, lag time and OD max for each strain were calculated using GrowthRates software with default parameters (Hall et al. 2014).

### Fermentation in beer wort and HPLC analysis

Fermentations were conducted using a 12°P high-gravity wort at 12°C in 50 mL (micro-fermentations). The 12 °P wort was prepared from a Munton’s Connoisseurs Pilsner Lager kit (Muntons plc, England). The worts were oxygenated to 15 mg/L prior to pitching. For the micro-fermentations, the strains were initially grown with constant agitation in 5 mL of wort for 48 hours at 15°C. Following this, 50 mL of fresh wort were inoculated to a final concentration of 15 × 10^6^ viable cells/mL and fermentations were maintained for seven days. Fermentations were weighed every day to calculate the CO_2_ output. The fermentations were maintained until no-CO_2_ lost was observed. At the end of the fermentation, the fermented worts were centrifuged at 9,000xg for 10 min and the supernatant was collected. From this, the concentration of extracellular metabolites was determined using HPLC. Specifically, 20 μL of filtered wort were injected in a Shimadzu Prominence HPLC (Shimadzu, USA) with a Bio-Rad HPX –87H column (Nissen et al., 1997). In this way, the concentrations of glucose, fructose, maltose, maltotriose, ethanol, and glycerol were estimated.

### Data Analysis

Multiple comparisons across localities were performed utilising a non-parametric Kruskal-Wallis test and Dunn’s Multiple Test Comparison. Genomewide *F*is and *F*st data across lineages was compared using paired Student t-test. Spearman rank correlation test and Pearson test were performed to determine correlations between variables. Finally, all analyses were performed utilising GraphPad Prism Software 5.2, except for correlation analysis which were performed in R (Development Core Team). In all cases *p*-values < 0.05 were considered as significant.

## ACKNOWLEDGMENTS

We would like to thank Valentina Abarca, Wladimir Mardones and Antonio Molina for technical help and Yessica Pérez, Antonia Nespolo, Natalia Hassan and Verónica Briceño for helping us in the sampling field trips recognising *Nothofagus* trees. We also thank Gilles Fischer for constructive feedback on the data analysis and the manuscript. We thank CONAF and WCS Chile for allowing us sampling yeasts across the country. This research was supported to FC by Comisión Nacional de Investigación Científica y Tecnológica CONICYT FONDECYT [1180161] and Millennium Institute for Integrative Biology (iBio). CV is supported by CONICYT FONDECYT [grant 3170404]. RN is supported by FIC ‘Transferencia Levaduras Nativas para Cerveza Artesanal’ and Fondecyt grant 1180917. K.U. was funded by USA 1899 – VRIDEI 021943CR-PAP Universidad de Santiago de Chile. Finally, we thank Ginkgobioworks for generating the sequence data.

## Authors Contributions

R.N. and F.C designed the study. R.N, F.C, C.V., C.O., S.T. collected and genotyped the strains. D.T. and E.M. performed the DNA sequencing. C.V, R.N., P.S. and F.C analysed the data. M.D.C., S.M. and G.L. performed the pangenome and FACS analysis. C.O., S.T., K.U., F.V. performed the experiments. R.N. and F.C wrote the manuscript.

---

**Figure S1. FACS analysis in *S. eubayanus isolates*. (A)** Fluorescence values for each sample are shown in grey. (*red): haploid CL609.1; (*green): diploid CL1004.1; (*blue): tetraploid CL1005.1. (B) Number of cell vs propidium iodide intensity is shown. Haploid (n), diploid (2n) and tetraploid (4n) examples are shown for the same strains as above (*).

**Figure S2. Results of STRUCTURE analysis for the different partition numbers (k = 3-7).** k-values were estimated for different partition numbers, being k = 5 the highest score.

**Figure S3. Principal Components Analysis using sequence similarity on SNP data.** A PCA analysis was performed using 229,272 SNPs. Strains clustered separately in agreement with the structure and phylogenetic analysis performed.

**Figure S4. Tajima D’ statistics of the PB-clades.** (A) Tajima D’ values along the genome for PB-1 (top), PB-2 (middle) and PB-3 (bottom) lineages. Tajima D’ were estimated using the R packages PopGenome 2.6.0. (B). Individual example of extremely low Tajima D’ values in Chromosome V for PB-1 and PB-2. The close-up denotes the genes located within the low Tajima D’ region suggesting a common genetic ancestry.

**Figure S5. Population differentiation values between lineages(*F***_**ST**_).

**Figure S6. Pairwise genetic distance between individuals versus geographic distance.** Genetic distances were estimated using the Nei’s distance method. Geographic distances were estimated based on map coordinates in google maps (https://www.google.com/maps). A positive correlation between genetic distance and geographic distance was found.

**Figure S7. Horizontal gene transfer event in PB-2 clade** (A) Nine ORFs have been identified on a single contig in 9 isolates. Around 6 kb of the flanking regions of these ORFs correspond to the chromosome I subtelomere, while the region where the ORFs are located do not show any similarities with known regions. (B) In the aminoacidic sequence of the nine ORFs, several domains can be identified of inferred by homologies.

**Figure S8. Principal Component Analysis of growth rates obtained under eight different conditions across isolates.**

**Figure S9. Agt1p analysis in *S. eubayanus*. (A).** Phylogeny of the *AGT1* gene on Chilean *eubayanus* strains. *AGT1_V, AGT1_XIII* and *AGT1_*XVI. Unrooted Neighbour-joining tree for the 82 Chilean strains and CBS12357/F1318. The trees were built using 29 SNPs and 6 INDELs for *AGT1_V*, 52 SNPs and 5 INDELs for *AGT1_XIII* and 6 SNPs and 2 INDELs for *AGT1_XVI.* (B). Modelling of the putative Agt1 structure in Chilean *S. eubayanus* strains. Three dimensional structure of the Agt1 transporter codified by *AGT1_V, AGT1_XIII* and *AGT1_XVI* in rainbow colouration (blue to red) from the N-terminus to the C-terminus. The truncated proteins consist of the first 50, 180 and 290 aminacids approximately for *AGT1_V, AGT1_XIII* and *AGT1_XVI* respectively. The truncated proteins are translated from the N-terminus to a premature STOP codon generated by INDELs on the sequence of the *AGT1* gene that codify the protein. (C) Binding affinity between the Agt1 transporter and maltose or maltotriose. Comparison between the binding affinities of the non-truncated and truncated Agt1 proteins with maltose. The binding affinities are predicted in Kcal/mol. Lower energy binding affinity (Kcal/mol) implies a higher affinity of the protein for the ligand. The binding affinity of all the truncated transporter are 32%, 14% and 7% lower than the non-truncated form for Agt1_V, Agt1_XIII and Agt1_XVI respectively (*** p-value < 0.001, **** p-value < 0.0001)

**Table S1. Number of samples obtained from each National Park.**

**Table S2. Bioinformatics Summary statistics together with NCBI accession numbers. Table S3. Population genetics summary statistics for each clade & locality.**

**Table S4. *F*st values per lineages and localities.**

**Table S5. Phenotype data for *S. eubayanus* strains.** (A). Raw phenotypic values. (B) Phenotype’s correlations

**Table S6. Phenotype comparison across localities.**

**Table S7. Fermentation data for *S. eubayanus* strains.** (A) CO_2_ lost in all strains. (B) Dunn’s multiple comparisons test across localities.

**Table S8. Physical Chemical parameters after wort fermentation.**

